## Appendix for "Habitat loss weakens the positive relationship between grassland plant richness and above-ground biomass"

Figure A1-A2

Table A1-A4

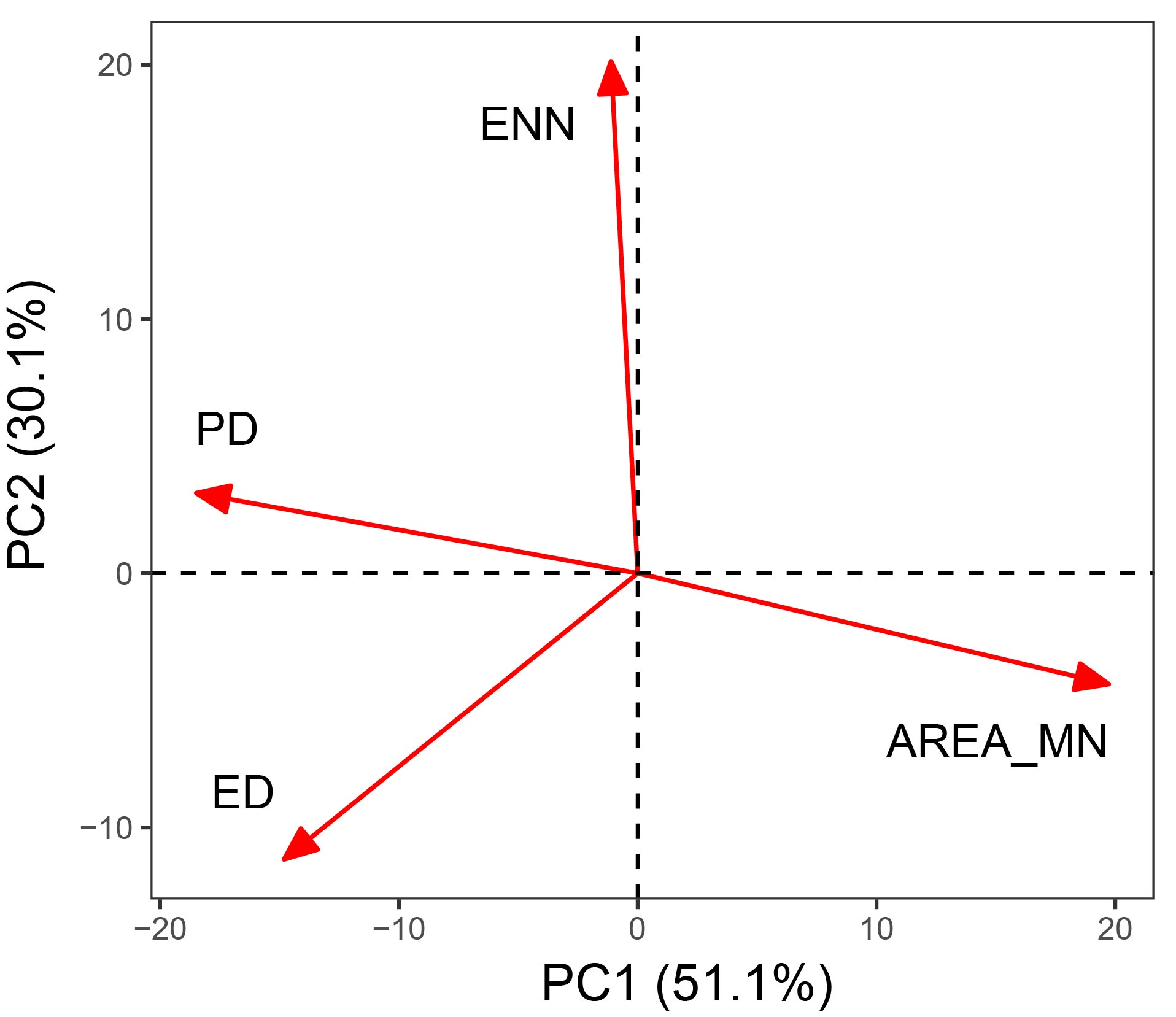

**Figure A1.** Biplot of the principal component analysis for calculating the single fragmentation index. PD: patch density; ED: edge density; AREA_MN: mean patch area; ENN: mean nearest-neighbor distance. PC1: the first principal component of the four fragmentation indices explained 51.1% of the variation. PC2: the second principal component of the four fragmentation indices explained 30.1% of the variation.

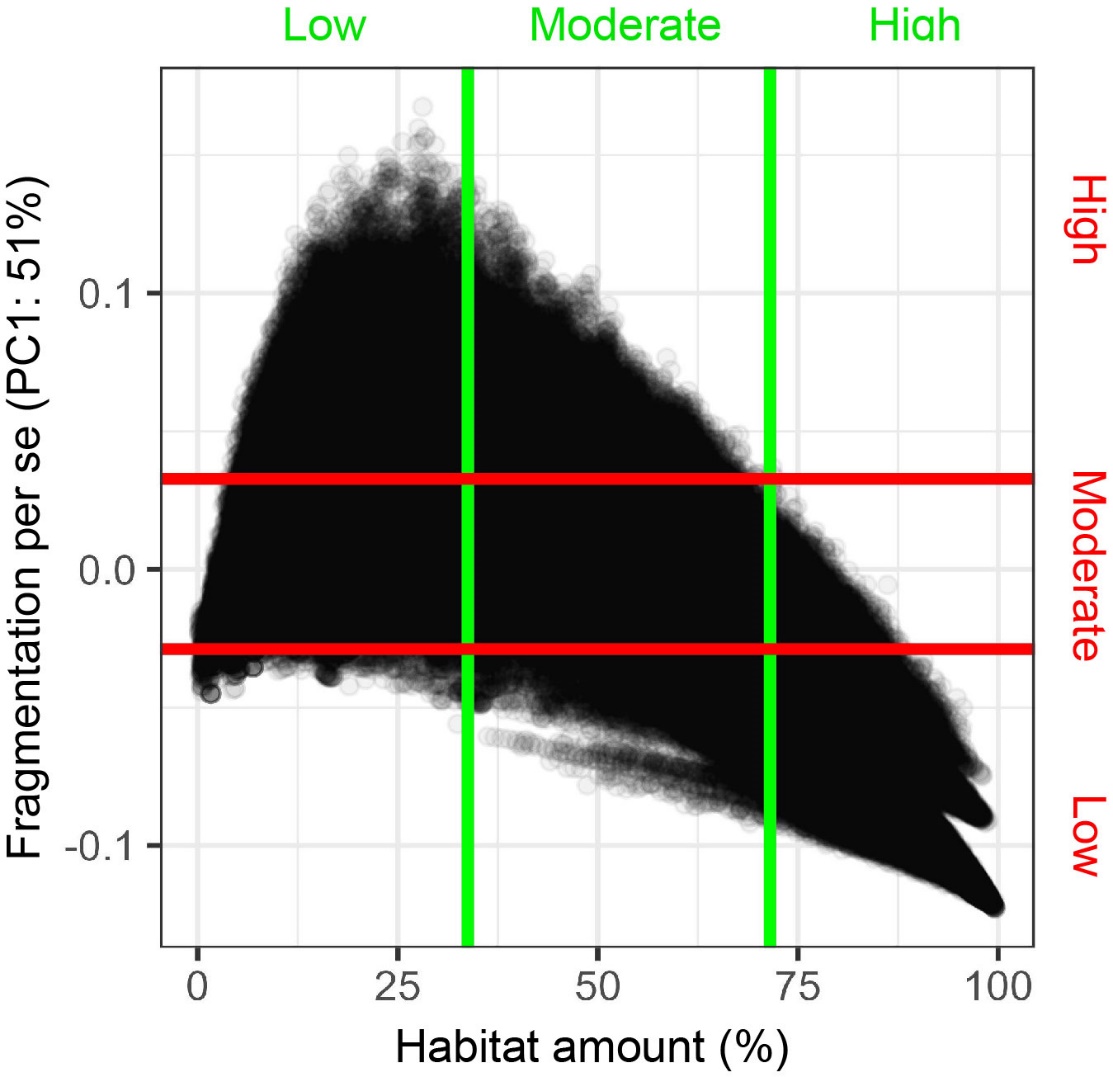

**Figure A2.** Scatterplot of habitat amount and fragmentation for stratified sampling. Each point in the figure represents the landscapes with 500-m radius. PC1: the first principal component of the four fragmentation indices. The green lines represent quartiles of habitat amount. The red lines represent quartiles of fragmentation per se. The high, moderate, and low on the horizontal axis represent the levels of habitat amount. The high, moderate, and low on the vertical axis represent the levels of fragmentation per se.

**Table A1****.** Summary of the principal component analysis for the four fragmentation indices.

| Results | PC1 |
| --- | --- |
| Eigenvalue | 2.04 |
| Proportion explained | 0.51 |
| Coefficient of the PD | -0.87 |
| Coefficient of the ED | -0.61 |
| Coefficient of the AREA_MN | 0.93 |
| Coefficient of the ENN | -0.17 |

Note: PD: patch density; ED: edge density; AREA_MN: mean patch area; ENN: mean nearest-neighbor distance; PC1: the first principal component of the four fragmentation indices.

**Table A2.** Variance inflation factors of predictor variables for above-ground biomass

| Variable | Variance inflation factor |
| --- | --- |
| Habitat loss | 2.49 |
| Fragmentation per se | 2.09 |
| Grassland specialist richness | 1.30 |
| Weed richness | 1.15 |
| Soil water content | 1.09 |
| Land surface temperature | 1.29 |

**Table A3.** Four optimal models of landscape context, environment factors, and plant diversity affecting above-ground biomass.

| Ranking | Model | R^2^ | AICc |
| --- | --- | --- | --- |
| 1 | AGB~HL+SWT+GSR+WR | 0.47** | 1213.7 |
| 2 | AGB~HL+FPS+SWT+GSR+WR | 0.47** | 1215.2 |
| 3 | AGB~HL+LST+SWT+GSR+WR | 0.47** | 1215.3 |
| 4 | AGB~FPS+SWT+GSR+WR | 0.46** | 1215.3 |

Note: AGB: above-ground biomass; HL: habitat loss; FPS: fragmentation per se; SWT: soil water content; LST: land surface temperature; GSR: grassland specialist richness; WR: weed richness; **: significance at the 0.01 level.

**Table A4.** List of 130 species of vascular plants recorded across 130 sites in this study.

| Family | Genus | Species | Habitat specialisation |
| --- | --- | --- | --- |
| Amaranthaceae | *Amaranthus* | *Amaranthus blitum* | W |
| Amaranthaceae | *Chenopodium* | *Chenopodium aristatum* | W |
| Amaranthaceae | *Bassia* | *Bassia scoparia* | W |
| Amaranthaceae | *Chenopodium* | *Chenopodium glaucum* | W |
| Amaranthaceae | *Suaeda* | *Suaeda glauca* | S |
| Amaranthaceae | *Corispermum* | *Corispermum mongolicum* | S |
| Amaranthaceae | *Kochia* | *Kochia prostrata* | S |
| Amaranthaceae | *Bassia* | *Bassia dasyphylla* | S |
| Amaranthaceae | *Corispermum* | *Corispermum chinganicum* | S |
| Amaranthaceae | *Salsola* | *Salsola collina* | W |
| Amaryllidaceae | *Allium* | *Allium anisopodium* | S |
| Amaryllidaceae | *Allium* | *Allium polyrhizum* | S |
| Amaryllidaceae | *Allium* | *Allium mongolicum* | S |
| Amaryllidaceae | *Allium* | *Allium bidentatum* | S |
| Amaryllidaceae | *Allium* | *Allium tenuissimum* | S |
| Amaryllidaceae | *Allium* | *Allium ramosum* | S |
| Apiaceae | *Bupleurum* | *Bupleurum scorzonerifolium* | S |
| Apiaceae | *Ferula* | *Ferula bungeana* | S |
| Apocynaceae | *Cynanchum* | *Cynanchum thesioides* | W |
| Asparagaceae | *Asparagus* | *Asparagus cochinchinensis* | S |
| Asteraceae | *Heteropappus* | *Heteropappus altaicus* | S |
| Asteraceae | *Artemisia* | *Artemisia argyi* | S |
| Asteraceae | *Xanthium* | *Xanthium sibiricum* | W |
| Asteraceae | *Saussurea* | *Saussurea amara* | S |
| Asteraceae | *Saussurea* | *Saussurea japonica* | S |
| Asteraceae | *Artemisia* | *Artemisia annua* | W |
| Asteraceae | *Leontopodium* | *Leontopodium leontopodioides* | S |
| Asteraceae | *Cirsium* | *Cirsium japonicum* | W |
| Asteraceae | *Ixeris* | *Ixeris polycephala* | W |
| Asteraceae | *Echinops* | *Echinops sphaerocephalus* | S |
| Asteraceae | *Artemisia* | *Artemisia frigida* | S |
| Asteraceae | *Serratula* | *Serratula centauroides* | S |
| Asteraceae | *Hemistepta* | *Hemistepta lyrata* | W |
| Asteraceae | *Hippolytia* | *Hippolytia trifida* | S |
| Asteraceae | *Taraxacum* | *Taraxacum mongolicum* | W |
| Asteraceae | *Echinops* | *Echinops gmelini* | S |
| Asteraceae | *Olgaea* | *Olgaea lomonossowii* | S |
| Asteraceae | *Ixeris* | *Ixeris gracilis* | W |
| Asteraceae | *Ixeridium* | *Ixeridium gracile* | W |
| Asteraceae | *Scorzonera* | *Scorzonera austriaca* | S |
| Asteraceae | *Artemisia* | *Artemisia halodendron* | S |
| Asteraceae | *Artemisia* | *Artemisia capillaris* | W |
| Asteraceae | *Neopallasia* | *Neopallasia pectinata* | S |
| Asteraceae | *Artemisia* | *Artemisia scoparia* | W |
| Asteraceae | *Filifolium* | *Filifolium sibiricum* | S |
| Boraginaceae | *Lappula* | *Lappula myosotis* | S |
| Brassicaceae | *Lepidium* | *Lepidium apetalum* | W |
| Brassicaceae | *Dontostemon* | *Dontostemon dentatus* | S |
| Brassicaceae | *Ptilotricum* | *Ptilotricum canescens* | S |
| Caryophyllaceae | *Gypsophila* | *Gypsophila davurica* | S |
| Caryophyllaceae | *Gypsophila* | *Gypsophila desertorum* | S |
| Caryophyllaceae | *Silene* | *Silene conoidea* | W |
| Caryophyllaceae | *Silene* | *Silene aprica* | S |
| Caryophyllaceae | *Gypsophila* | *Gypsophila licentiana* | S |
| Convolvulaceae | *Convolvulus* | *Convolvulus arvensis* | W |
| Convolvulaceae | *Convolvulus* | *Convolvulus ammannii* | S |
| Crassulaceae | *Orostachys* | *Orostachys fimbriatus* | S |
| Cyperaceae | *Carex* | *Carex korshinskyi* | S |
| Ephedraceae | *Ephedra* | *Ephedra sinica* | S |
| Euphorbiaceae | *Euphorbia* | *Euphorbia humifusa* | W |
| Euphorbiaceae | *Euphorbia* | *Euphorbia esula* | S |
| Fabaceae | *Astragalus* | *Astragalus scaberrimus* | S |
| Fabaceae | *Astragalus* | *Astragalus melilotoides* | S |
| Fabaceae | *Astragalus* | *Astragalus dahuricus* | S |
| Fabaceae | *Oxytropis* | *Oxytropis bicolor* | S |
| Fabaceae | *Oxytropis* | *Oxytropis diversifolia* | S |
| Fabaceae | *Lespedeza* | *Lespedeza bicolor* | S |
| Fabaceae | *Medicago* | *Medicago ruthenica* | S |
| Fabaceae | *Corethrodendron* | *Corethrodendron fruticosum var. mongolicum* | S |
| Fabaceae | *Gueldenstaedtia* | *Gueldenstaedtia verna* | S |
| Fabaceae | *Thermopsis* | *Thermopsis lanceolata* | S |
| Fabaceae | *Astragalus* | *Astragalus galactites* | S |
| Fabaceae | *Oxytropis* | *Oxytropis racemosa* | S |
| Fabaceae | *Oxytropis* | *Oxytropis leptophylla var. turbinata* | S |
| Fabaceae | *Astragalus* | *Astragalus adsurgens* | S |
| Fabaceae | *Lespedeza* | *Lespedeza daurica* | S |
| Gentianaceae | *Gentiana* | *Gentiana dahurica* | S |
| Gentianaceae | *Gentiana* | *Gentiana scabra* | S |
| Geraniaceae | *Erodium* | *Erodium stephanianum* | S |
| Iridaceae | *Iris* | *Iris lactea* | S |
| Iridaceae | *Iris* | *Iris lactea var. chinensis* | S |
| Iridaceae | *Iris* | *Iris tenuifolia* | S |
| Lamiaceae | *Dracocephalum* | *Dracocephalum heterophyllum* | S |
| Lamiaceae | *Thymus* | *Thymus mongolicus* | S |
| Lamiaceae | *Scutellaria* | *Scutellaria scordifolia* | S |
| Lamiaceae | *Phlomis* | *Phlomis umbrosa* | S |
| Lamiaceae | *Lagochilus* | *Lagochilus ilicifolius* | S |
| Lamiaceae | *Dracocephalum* | *Dracocephalum moldavica* | S |
| Lamiaceae | *Lagopsis* | *Lagopsis supina* | W |
| Linaceae | *Linum* | *Linum usitatissimum* | W |
| Nitrariaceae | *Peganum* | *Peganum harmala* | S |
| Orobanchaceae | *Cymbaria* | *Cymbaria dahurica* | S |
| Plantaginaceae | *Plantago* | *Plantago asiatica* | W |
| Plantaginaceae | *Plantago* | *Plantago depressa* | W |
| Plumbaginaceae | *Limonium* | *Limonium aureum* | S |
| Poaceae | *Echinochloa* | *Echinochloa crusgali* | W |
| Poaceae | *Cleistogenes* | *Cleistogenes hancei* | S |
| Poaceae | *Agropyron* | *Agropyron cristatum* | S |
| Poaceae | *Cleistogenes* | *Cleistogenes squarrosa* | S |
| Poaceae | *Stipa* | *Stipa breviflora* | S |
| Poaceae | *Setaria* | *Setaria viridis* | W |
| Poaceae | *Eragrostis* | *Eragrostis pilosa* | W |
| Poaceae | *Enneapogon* | *Enneapogon borealis* | S |
| Poaceae | *Alopecurus* | *Alopecurus aequalis* | W |
| Poaceae | *Phragmites* | *Phragmites australis* | W |
| Poaceae | *Elymus* | *Elymus dahuricus* | S |
| Poaceae | *Koeleria* | *Koeleria cristata* | S |
| Poaceae | *Bromus* | *Bromus inermis* | W |
| Poaceae | *Cleistogenes* | *Cleistogenes songorica* | S |
| Poaceae | *Leymus* | *Leymus chinensis* | S |
| Poaceae | *Stipa* | *Stipa capillata* | S |
| Poaceae | *Digitaria* | *Digitaria ischaemum* | W |
| Poaceae | *Stipa* | *Stipa grandis* | S |
| Polygonaceae | *Polygonum* | *Polygonum aviculare* | W |
| Polygonaceae | *Polygonum* | *Polygonum sibiricum* | W |
| Portulacaceae | *Portulaca* | *Portulaca oleracea* | W |
| Primulaceae | *Androsace* | *Androsace umbellata* | S |
| Primulaceae | *Lysimachia* | *Lysimachia barystachys* | W |
| Ranunculaceae | *Thalictrum* | *Thalictrum petaloideum* | S |
| Ranunculaceae | *Thalictrum* | *Thalictrum squarrosum* | S |
| Rosaceae | *Potentilla* | *Potentilla betonicifolia* | S |
| Rosaceae | *Chamaerhodos* | *Chamaerhodos erecta* | S |
| Rosaceae | *Potentilla* | *Potentilla bifurca* | W |
| Rosaceae | *Potentilla* | *Potentilla anserina* | W |
| Rosaceae | *Potentilla* | *Potentilla verticillaris* | S |
| Rosaceae | *Geum* | *Geum aleppicum* | W |
| Rosaceae | *Potentilla* | *Potentilla acaulis* | S |
| Rutaceae | *Haplophyllum* | *Haplophyllum dauricum* | S |
| Thymelaeaceae | *Stellera* | *Stellera chamaejasme* | S |
| Zygophyllaceae | *Tribulus* | *Tribulus terrester* | W |

Note: S: Grassland specialists; W: Weeds.
